# Sedative choice alters *Klebsiella pneumoniae* lung pathogenesis and dissemination

**DOI:** 10.1101/2025.05.27.656367

**Authors:** David Mains, Ella R. Rotman, Deanna K. Aman, Denise A. Ludvik, Anirudh Desikan, Acadia A. Kocher, An N.H. Tran, Mark J. Mandel, Nancy E. Freitag

**Affiliations:** Department of Microbiology and Immunology, University of Illinois at Chicago, Chicago, IL 60612; Department of Microbiology-Immunology, Northwestern University Feinberg School of Medicine, Chicago, IL 60611; Department of Pharmaceutical Sciences, University of Illinois, Chicago, IL 60611; Department of Medical Microbiology and Immunology, University of Wisconsin-Madison, Madison, WI 53706

**Keywords:** *Klebsiella pneumoniae*, transposon insertion sequencing (INSeq/Tn-seq), pneumonia, propofol, ketamine, virulence

## Abstract

*Klebsiella pneumoniae* make up 85% of carbapenem-resistant Enterobacteriaceae (CRE), bacteria that have become an urgent threat to public health. *K. pneumoniae* is largely transmitted in healthcare settings, where inpatients and outpatients are often anesthetized with the widely-used anesthetic induction agent propofol. Recent evidence obtained from rodent infection models indicates that propofol exposure can dramatically increase host susceptibility to microbial infections. Given that intensive care patients who are at a greater risk for *K. pneumoniae* lung infections are often given propofol during their hospitalization, we investigated the outcome of *K. pneumoniae* infections in mice briefly sedated with either propofol or ketamine/xylazine as control. Propofol-sedated mice experienced more rapid dissemination from the lungs to secondary sites of infection and their lungs exhibited more severe pathology. Based on these observations, we investigated bacterial factors involved in infection and dissemination in mice with propofol or ketamine/xylazine sedation using a high throughput insertion sequencing (INSeq) approach. We identified numerous novel potential virulence factors together with previously identified gene products, confirming the validity of our screen. We further characterized a mutant lacking the phospholipid retrograde trafficking chaperone MlaC and found that the degree of mutant attenuation was dependent upon sedation method. These results highlight the importance of sedative choice when studying hospital-acquired microbial infections and suggest that sedation can influence outcome of *K. pneumoniae* infection and dissemination in animal models.

**IMPORTANCE:** Host sedation by either propofol or ketamine exposure differentially impacted the severity of *K. pneumoniae* lung infection following intranasal inoculation of mice. While propofol-sedated mice exhibited increased lung pathology, some bacterial mutants, such as those lacking the MlaC gene product associated with the maintenance of inner and outer membrane lipid asymmetry, exhibited more severe attenuation following propofol sedation versus ketamine/xylazine. Given the dominating use of propofol in health care settings for the induction and maintenance of anesthesia, procedural sedation, and sedation for intensive care patients, this work provides important perspective as to how the choice of an anesthetic agent may impact the outcome of healthcare-associated infections.

## INTRODUCTION

Healthcare-associated infections (HAIs) pose the threat of increased morbidity for many patients while serving as a major cause of mortality for intensive care patients. The anesthetic induction agent propofol is widely used in critical care settings and is a common choice for light sedation in the treatment for critically ill patients (1). Furthermore, patients requiring mechanical ventilation are frequently administered sedatives (2) and are at increased risk for a variety of complications including pneumonia, collectively referred to as ventilator-associated events (VAEs) (3–6). The recent global pandemic of COVID-19, caused by SARS-CoV-2, has highlighted risks of ventilator treatment (7–11). Mechanical ventilation increases the risk of aspirating commensal bacteria into the lungs, and use of sedation during ventilation is associated with an increased hazards ratio for VAEs (5–6).

The general anesthetic propofol has grown to become the predominant intravenous anesthetic in clinical use throughout the world, lauded for its rapid onset and offset of sedation, relatively mild side effects, and flexibility in both the induction and maintenance of anesthesia (12, 13). While there have been multiple reports on propofol’s immunomodulatory effects *in vitro* (13–24), *in vivo* evidence has only recently begun to be reported. Klompas et al. correlated propofol sedation in mechanically-ventilated patients with a significantly increased risk of VAEs, including pneumonia, compared to patients not receiving propofol (5). Visvabharathy et al. reported that acute propofol sedation dramatically increased susceptibility to both *Listeria monocytogenes* and *Staphylococcus aureus* bloodstream infections in a mouse model and additionally led to more severe pathology at primary sites of infection (25, 26). This susceptibility correlated with a reduced capacity to effectively recruit mature immune effector cells to sites of infection following propofol sedation. These propofol-associated influences on host immunity are especially important to consider given that propofol is administered widely in both inpatient and outpatient procedures and suggested to us that propofol could have a wider impact beyond the intravenously administered gram-positive pathogens we had previously examined.

To explore the potential impact of propofol sedation on the outcome of respiratory infections, we have made use of a mouse model of lung infection with the opportunistic pathogen *Klebsiella pneumoniae*. *K. pneumoniae* is a gram-negative extracellular bacterium that is ubiquitous in the water and soil of the environment and also colonizes the nasopharynx and intestinal tract of humans and animals (27). It accounts for 3-8% of all nosocomial bacterial infections including pneumonia, gram-negative bacteremia, and urinary tract infections (28). *K. pneumoniae* is the second most common cause of gram-negative bacteremia in the United States with 12-50% of *K. pneumoniae* bloodstream infections originating from the lungs (28–31). A significantly worse outcome of bacteremia is associated with *K pneumoniae* infections from a respiratory source (29). *K. pneumoniae* is a growing threat worldwide as a lethal nosocomial pathogen that is rapidly acquiring antimicrobial resistance (32).

Using an intranasal model of *K pneumoniae* infection in mice, we compared the impact of propofol versus ketamine/xylazine sedation on bacterial colonization of the lung, lung pathology, and the dissemination of bacteria to distal organs. We further investigated whether sedative choice influenced the requirement for distinct *K pneumoniae* virulence determinants using a whole genome transposon insertion sequencing (INSeq) approach.

## MATERIALS AND METHODS

### Bacterial media and strains

*K. pneumoniae* and *Escherichia coli* strains used in this study are listed in **Table 1**. Strains were grown at 37 °C in lysogeny broth (LB; per liter, 10 g Bacto-tryptone [BD], 5 g yeast extract [BD], 10 g NaCl [Sigma-Aldrich]). When cells were grown in the presence of hygromycin, low-salt LB was used (i.e., 5 g NaCl per liter). When necessary, antibiotics (Gold Biotechnology) were added to the media at the following concentrations: rifampicin 30 μg ml^-1^, hygromycin 100 µg ml^-1^, kanamycin 50 µg ml^-1^, and carbenicillin 100 µg ml^-1^. When necessary, the following compounds were added to the media to the indicated final concentrations: sodium salicylate (Sigma-Aldrich): 2 mM; diaminopimelate (DAP; Sigma-Aldrich): 0.3 mM; arabinose (Gold Biotechnology): 100 mM. Growth media were solidified with 1.5% agar as needed. Standard molecular biology techniques were used to introduce plasmids into *E. coli* (i.e. electroporation) and into *K. pneumoniae* (i.e., electroporation or conjugation).

**Table 1.**
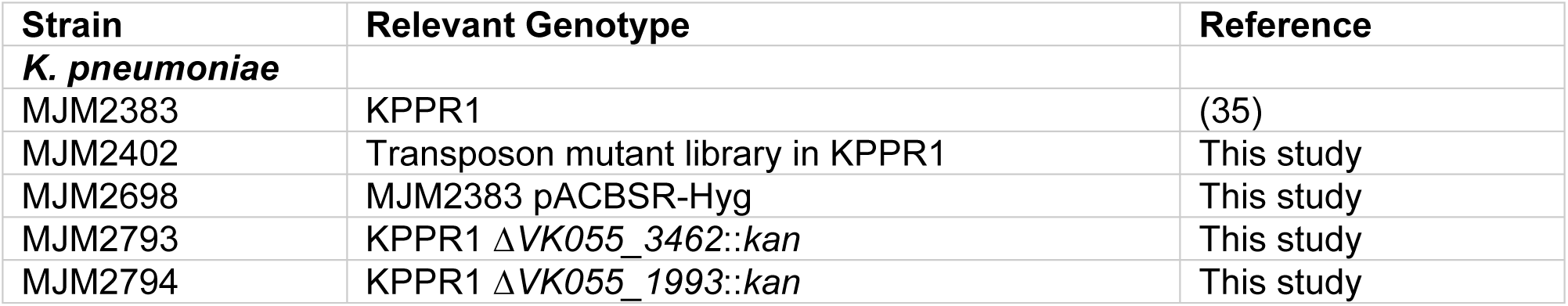
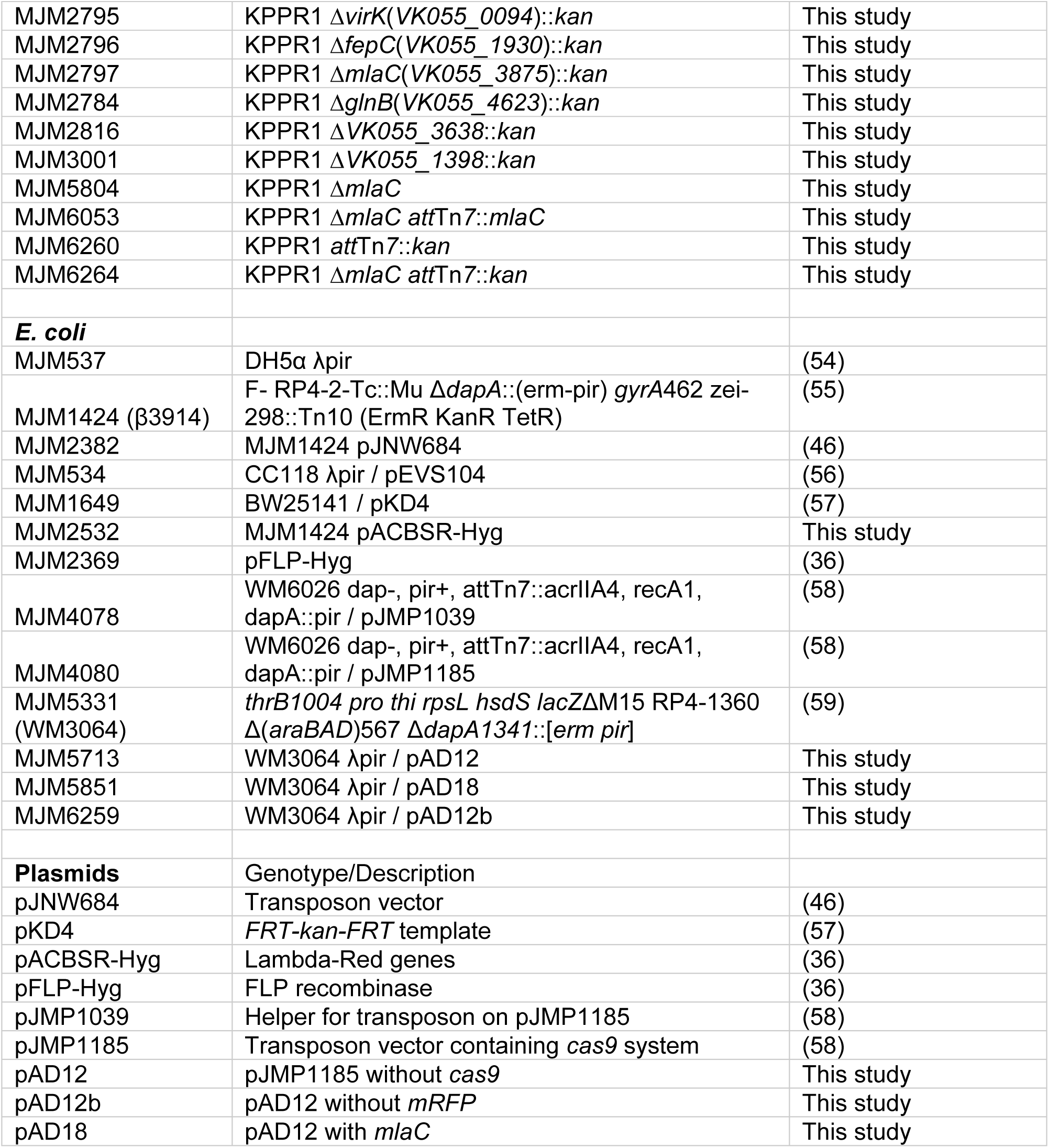
Bacterial strain and plasmid list.

### Animal infections

All animal procedures were approved by the University of Illinois Chicago Animal Care Committee and were conducted in the Biological Resources Laboratory. *K. pneumoniae* strains were cultured overnight with shaking (37°C, 180 rpm) in LB broth. The following morning, strains were subcultured 1:50 in fresh LB and incubated with shaking for one hour to an OD_600_ of 0.2-0.4. Optical density was adjusted to 0.2 OD_600_ with LB broth and bacteria were recovered by centrifugation, washed, and diluted to the desired CFU in sterile 1X phosphate-buffered saline (PBS) for infection. Strains were kept on ice until immediately before infection. Animals were sedated via tail vein injection with ketamine (20 mg/kg; Zoetis, Kalamazoo, MI) and xylazine (4 mg/kg; Akorn, Decatur, IL) in Intralipid as control or propofol (20 mg/kg; Zoetis) in Intralipid. For non-competitive infections, approximately 2.5 x 10^2^ colony forming units (for propofol sedated mice) or 2.5 x 10^3^ colony forming units (ketamine/xylazine sedated mice) of each strain was administered via intranasal route into 8- to 10-week old Swiss Webster mice (Envigo, Madison, WI) and for competitive infections, approximately 5 x 10^3^ CFU of WT and a mutant were premixed and administered via intranasal route for a total 1 x 10^4^ CFU dose. For non-competitive infections, mice were sacrificed after 24 or 48 hours. Lungs, livers, and spleens were isolated, homogenized, and plated in triplicate on LB agar containing rifampicin for enumeration of bacterial burdens in each organ. For competitive infections, mice were sacrificed after 24 hours and lungs were homogenized and plated in triplicate on LB agar containing rifampicin and on LB agar containing rifampicin and kanamycin.

### Histology

For histological analysis, mice were sacrificed individually via CO_2_ asphyxiation. Lungs were inflated with PBS at a flow rate no greater than 200 μl/second using a 3 ml syringe with a 22 g needle inserted into the trachea. Once lungs were fully inflated and prior to removal of the needle, the trachea was tied off with a surgeon’s knot of suture thread. Lungs were placed in 30 ml of neutral buffered formalin for 48 hours, then transferred into 70% ethanol. Paraffin-embedded sections were prepared by the UIC Research Histology and Tissue Imaging core facility. Five micron-thick sections were stained with hematoxylin and eosin (H&E) and mounted. Digital scans of H&E-stained lung sections were captured on a Keyence digital microscope and analyzed using ImageJ software.

### H&E histopathological analyses

Mouse lung images standardized by containment of only the left lobe and cranial and caudal portions of the right lobe were analyzed on ImageJ software by an examiner blinded to sedative treatment. Total lesion area was quantified by the sum of basophilic areas of inflammatory cell infiltration as a fraction of total lung area per image. Number or lesions and average legion size composing the total lesion area were also enumerated per whole lung slice.

### Immunofluorescence

For immunofluorescent analysis, mice were sacrificed via isoflurane. The lung vasculature was perfused with 10 ml ice cold PBS containing 50U/ml heparin to remove blood and lungs were infused with one milliliter of neutral buffered formalin solution (Millipore Sigma). Lungs were placed in 20 ml of neutral buffered formalin for 48 hours, then transferred into 70% ethanol. Unstained, paraffin-embedded sections of 5μm thickness were prepared by the UIC Research Histology and Tissue Imaging core facility. Paraffin was dissolved in xylenes and sections were rehydrated in graded ethanol washes, followed by washing in tris-buffered saline (TBS, pH 7.6). Antigen retrieval was carried out in 10 mM sodium citrate buffer with 0.05% Tween-20 (pH 6.0) in a pressure cooker on high temperature for fifteen minutes. Slides were blocked with Background Buster (Innovex, Richmond, CA). Primary antibody was *K. pneumoniae* rabbit polyclonal (ThermoFisher, Waltham, MA) and secondary antibody was goat anti-rabbit-FITC (Invitrogen, Carlsbad, CA). Slides were mounted with ProLong Gold antifade with DAPI (ThermoFisher), imaged using a Zeiss Axio Imager A2 upright microscope (Carl Zeiss, Thornwood, NY), and analyzed on Zen 2012 (Carl Zeiss) imaging software.

### Generation of KPPR1 transposon mutant library

KPPR1 and MJM2382 were grown overnight in LB and LB carb DAP, respectively, at 37 °C with aeration. Saturated cultures were diluted 1:80 in 3 ml and grown for 2 hours and 40 minutes at 37 °C. 2.5 ml of *E. coli* and 5 ml of *K. pneumoniae* were pelleted and combined in 500 µl LB, which was spotted in fifty 10-µl spots on LB. After 3.5 hours at 37 °C, each spot was swabbed into 500 µl LB, vortexed, and 50 µl was spread onto LB plates containing kanamycin and salicylic acid for overnight incubation at 37 °C (the presence of salicylic acid reduces the amount of capsule produced). An estimated 50,000 colonies were swabbed into 25 ml LB mixed with 12.5 ml 50% glycerol on ice and frozen at −80°C with a final density of 3 x 10^11^ CFU/ml.

### INSeq analysis

Genomic DNA was extracted using a Maxwell 16 instrument (Promega, Madison, WI). Transposon insertion junctions were amplified from three input samples and six output samples according to the previously published protocol (33). Samples were single-end (SE50) sequenced with an Illumina HiSeq 2500 platform (Illumina, San Diego, CA) at the Tufts University Genomics Core Facility in Boston, MA with 45 million reads generated. FASTQ files were first trimmed to the chromosomal sequence flanking each transposon insertion with pyinseq v0.2 (https://github.com/mjmlab/pyinseq). Files were subsequently analyzed either with the full pyinseq v0.2 pipeline to obtain cpm (normalized counts-per-million) data for each replicate or with TRANSIT 3.8 (34) using the ‘resampling’ algorithm against the KPPR1 genome sequence CP009208.1 (35) with parameters -a -iN 5 -iC 10. Sequencing data were deposited at NCBI SRA under accessions SRR6480865-SRR6480873 as detailed in **Table S1**. A table of pyinseq analyzed values for each KPPR1 gene is posted as **Table S2**. Significant results from the TRANSIT output are listed in **Table S3** and **Table S4**.

### Generation of *K. pneumoniae* gene deletion strains

Deletions were made in the KPPR1 background using Lambda Red recombination (36). Knockout cassettes for each gene were generated using the “KO” primers listed in **Table S3** to amplify the KanR cassette from pKD4 with homologous ends to chromosomal location of each gene. KPPR1 was electroporated with pACBSR-Hyg to create strain MJM2698. This strain was electroporated with each of the knockout cassettes. Strains with Kan resistance were selected and then patched onto low salt-LB-Hyg plates to confirm the loss of pACBSR-Hyg. Kan-resistant and Hyg-sensitive isolates were verified with PCR using the verification (V) primers listed in **Table S5**. The PCR product was submitted for Sanger sequencing at the Northwestern University Sanger Sequencing Core to confirm the integrity of the junctions where the knockout cassette replaced the gene of interest. To remove the KanR cassette, strains were electroporated with pFLP-Hyg and acquisition of the plasmid was selected on low salt-LB-Hyg media. Candidates were incubated at 42 °C overnight and then screened for loss of Kan resistance. Colonies that lost Kan resistance were passaged for several days on LB plates and then screened for Hyg sensitivity to confirm loss of pFLP-Hyg. PCR was used to confirm removal of the Kan cassette using the verification (V) primers listed in **Table S5**.

### Construction of *K. pneumoniae* complementation strains

Complementation of deleted genes was accomplished by integrating wild-type alleles of the gene at the presumed neutral *att*Tn*7* chromosomal site using a site-specific Tn*7* transposon (**Figure S2**). Published vector pJMP1185 expresses a Cas9 system with a constitutive promoter driving expression of *mRFP* as a control (37). We first removed the *cas9* gene from the transposon to create pAD12. Primers AD_254 and AD_255 were used to amplify pJMP1185 without the *cas9* gene. The plasmid was ligated back together with T4 ligase and transformed into WM3064 *E. coli* cells. Candidates were selected on LB-Kan plates, plasmid insertion was verified with PCR using primers AD_191 and AD_192, and the correct sequence was confirmed with Oxford Nanopore sequencing (Plasmidsaurus). To generate complement vectors, the *mRFP* gene was replaced with one of the genes of interest to generate pAD14, pAD16, pAD18, or completely removed to generate an empty vector control (pAD12b). Primers AD_258 and AD_259 were used to amplify pAD12. The reaction was PCR purified and DpnI digested. Genes were amplified with either AD_264/AD_265, AD_272/AD_273, or AD_276/AD_277 and then cloned into the backbone using the NEB Hi-Fi assembly kit. For the empty vector control, primers DAT_535/DAT_536 were used to amplify the backbone and the Q5 site-directed mutagenesis kit (NEB) was used for assembly. For the complementation plasmids and empty vector control, the assembly mix was transformed into WM3064 *E. coli* cells and candidates were selected on LB-Kan. The resulting plasmids were verified with Oxford Nanopore sequencing (Plasmidsaurus). Plasmids were transformed into relevant *K. pneumoniae* strains using conjugation. Candidates were selected using Kan resistance and verified using PCR (DAT_529/DAT_530) and Oxford Nanopore sequencing (Plasmidsaurus).

### SDS/EDTA susceptibility

To assess doubling time, strains were grown in LB broth at 37°C with shaking for 3 h, with samples taken every 30 min. Serial dilutions were spot plated to enumerate viable CFU and the doubling time was calculated from these points. To assess growth on SDS/EDTA, cultures were grown to saturation in LB, serially diluted, and 5 μl was spotted on LB plates or LB plates containing 0.1% SDS/1.5 mM EDTA. Plates were incubated overnight at 37°C and imaged with a Nikon D810 digital camera.

### Statistical Analysis

For mouse experiments that compared multiple groups, bacterial burden data was log10 transformed and analyzed via the Kruskal Wallis test with Dunn’s post-hoc test. Data graphing and analysis was conducted on GraphPad Prism software.

## RESULTS

### Sedation with the anesthetic propofol leads to increased lung colonization with *K. pneumoniae*

Previous investigations into the impact of propofol on bloodstream infections using the gram-positive pathogens *L. monocytogenes* and *S. aureus* revealed significant alterations in bacterial colonization, growth, and pathology in target organs (25). Given the prevalence of *K. pneumoniae* acquisition in health-care settings where propofol is widely used, and to examine the impact of propofol on a model gram-negative pathogen, we assessed whether propofol affected lung infection of *K. pneumoniae* via an intra-nasal inoculation route. Female Swiss Webster mice were briefly sedated (5-10 minutes) with either intravenous propofol or intravenous ketamine/xylazine and intranasally infected with 3×10^4^ CFU KPPR1. Bacterial burdens were assessed at 24 and 48 hours post infection in the lung, liver, and spleen. Brief propofol sedation had no measurable impact on lung burdens 24 hours post-infection, but burdens were increased nearly 100-fold by 48 hours compared to ketamine/xylazine-sedated controls (**Figure 1**). We also assessed the impact on dissemination from the lungs and again found that while secondary organs had comparable burdens at 24 hours, average burdens in both livers and spleens at 48 hours were more than 1,000-fold higher than ketamine/xylazine sedated controls (**Figure 1**). We conclude that the increased susceptibility of murine hosts to infection observed following propofol sedation is therefore broad enough to impact multiple routes of infection with both gram-positive and gram-negative bacteria.

**Figure 1.**
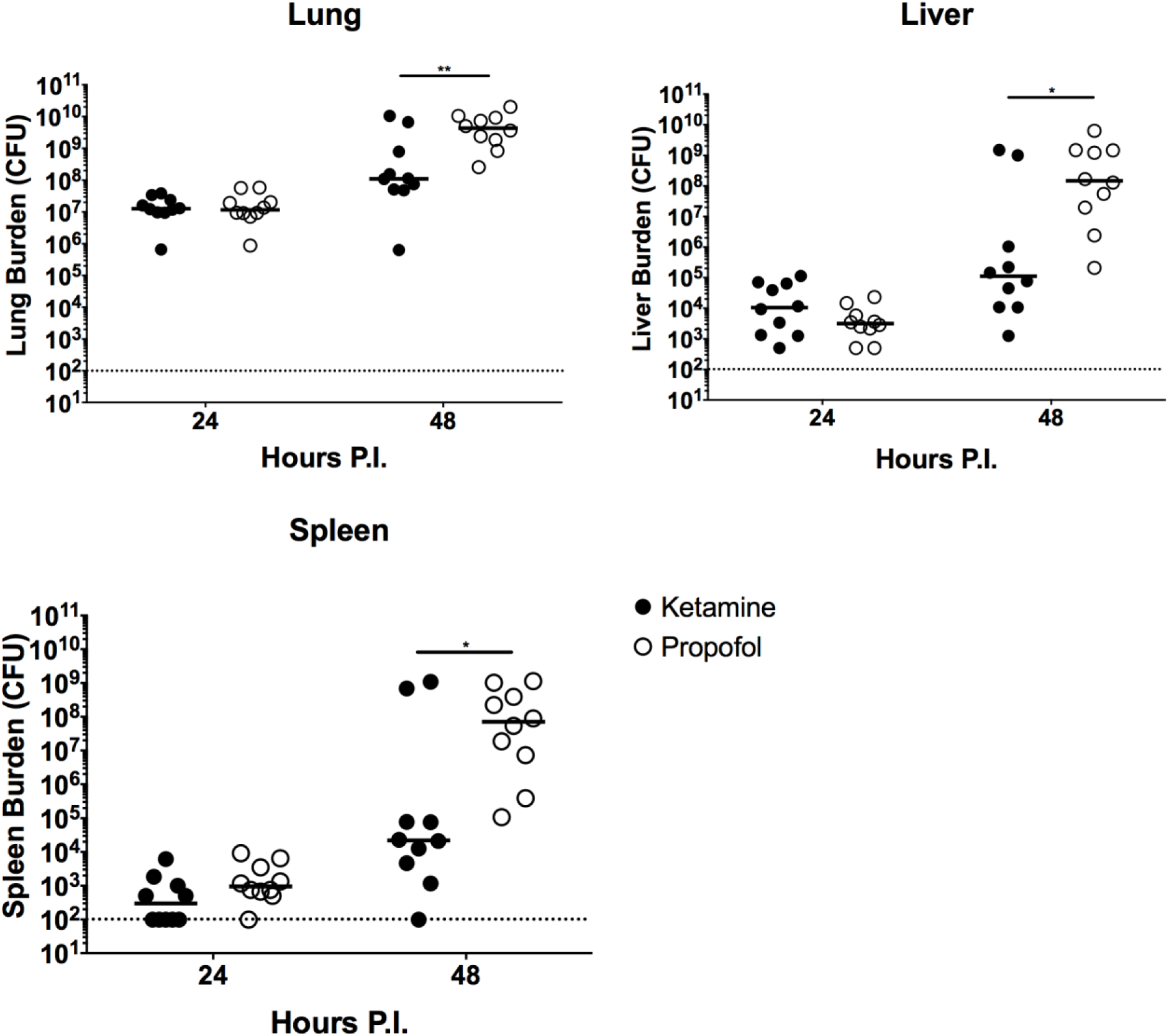
Brief propofol sedation increases *Klebsiella* lung burdens and dissemination to secondary tissues. Swiss Webster mice were sedated with either ketamine/xylazine (closed circles) or propofol (open circles) and infected with 3×10^4^ CFU intra-nasally and lung, liver, and spleen burdens were assessed at 24 and 48 hours post infection. Statistics were calculated using the Mann Whitney U test (*, p ≤ 0.05; **, p ≤ 0.01).

### Propofol exacerbates lung pathology associated with *K. pneumoniae* infection

Previous research using propofol sedation prior to infection identified gross changes in pathology at primary sites of infection (25, 26). When examining lung histology, we observed that by 48 hours post-infection severe damage was evident in lungs regardless of sedation agent (**Figure 2A-B**), Quantification of these regions showed that propofol-sedated mice contained significantly larger lung lesions while the total number of focal lesions did not differ based on sedative agent (**Figure 2D-F**). Notably, only propofol-sedated mouse lungs featured large areas of de-cellularized or denuded tissue containing little identifiable structure and only sporadic immune infiltration. Immunofluorescent examination of these regions revealed that *K. pneumoniae* had completely overrun local containment and appeared to have developed a replicative niche (**Figure 2G-H**). It is likely that these bacteria-packed regions underlie the disparate burdens observed between sedation groups.

**Figure 2.**
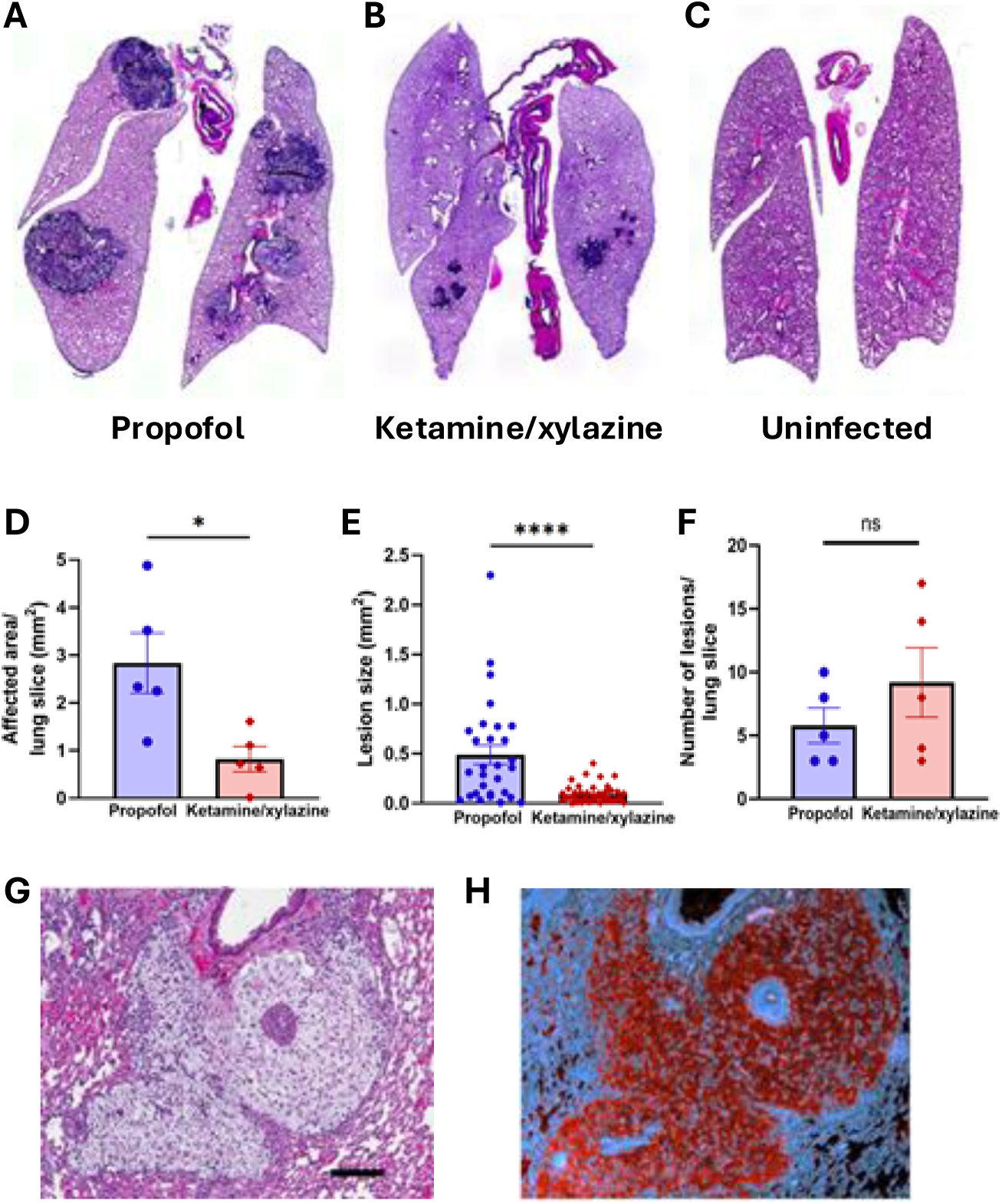
Histological analysis of *K. pneumoniae* lung pathology 48 hours post-infection. Swiss Webster female mice were infected with 1×10^2^ CFU KPPR1. Representative H&E images of *Kp* infected mouse whole lung tissue 48H post infection after propofol (A), ketamine/xylazine (B) sedation, or uninfected control (C). (D) Quantification of total lesioned area per lung section (n=5 per treatment). (E) Area of individual lesion sizes. (F) Quantification of number of lesions in whole lung sections from left and right (cranial and caudal) lung lobes (n=5 per treatment). (G-H) Representative H&E (G) and Immunofluorescent staining (H) of propofol-sedated mouse lungs with the propofol specific histopathological feature of acellular denuded regions packed with *K. pneumoniae*. Black bar represents 100µm. Statistics were calculated using an unpaired, two-tailed Student’s t-test (*, p ≤ 0.05; ****, p<.0001).

### Transposon insertion sequencing identifies *K. pneumoniae* genes that are required for outgrowth in the lung in a sedative-independent manner

Genome-wide approaches for the discovery of virulence factors required for *in vivo* growth have been applied in *K. pneumoniae* previously with considerable success (38–43). In addition to examining the impact of propofol on *K. pneumoniae* lung infection, we also sought to investigate what impact propofol might have on the *K. pneumoniae* global virulence repertoire required for successful growth in the lung. Transposon insertion sequencing (INSeq/Tn-seq) is a powerful, high resolution approach to analyze whole genomes by coupling Mariner transposon mutagenesis with high-throughput sequencing to rapidly and comprehensively quantify the relative fitness of large pools of transposon mutants (44–46). We applied this approach by mutagenizing *K. pneumoniae* strain KPPR1, sedating mice with either propofol or ketamine/xylazine, followed by the infection of mice via the intranasal route. We generated a 50,000 mutant transposon library in KPPR1 using the plasmid pJNW684 (46). Hits were observed across the chromosome with an average of approximately 9.1 insertions per kb. Three propofol-sedated and three ketamine/xylazine control-sedated mice were intranasally infected with the library at a dose of 6 x 10^5^ CFU for 24 hours, after which the lungs were harvested and genomic DNA was extracted. The 24 h time point was selected to identify mutants with fitness defects in advance of the lethality differences that begin to occur at 48 h with the use of a higher CFU dose and so that the analysis was not complicated by the high proportion of lethality at the later time point. The higher CFU dose of 6 x 10^5^ CFU was selected to increase mutant representation within the library for the subsequent INSeq analyses.

We analyzed the output data to examine genes that displayed fitness defects under both sedatives. Using the software TRANSIT (34), we identified 217 genes for which mutants exhibited significant depletion (and 2 genes for which mutants exhibited enrichment) under both sedative conditions (**Fig. 3**; **Tables S3, S4**). The depleted mutants match factors identified in previous transposon insertion sequencing studies, including *aroA, copA, hisA, ilvC, leuA, leuB, serA, serB, trpE,* and *tyrA* (Fig. 3) (43, 47). Identification of conserved responses across the two sedatives, and identification of responses consistent with published studies supports the quality of the INSeq analysis and indicates that targeting of these pathways is likely to yield benefits to the extent that propofol responses are conserved in human patients.

**Figure 3:**
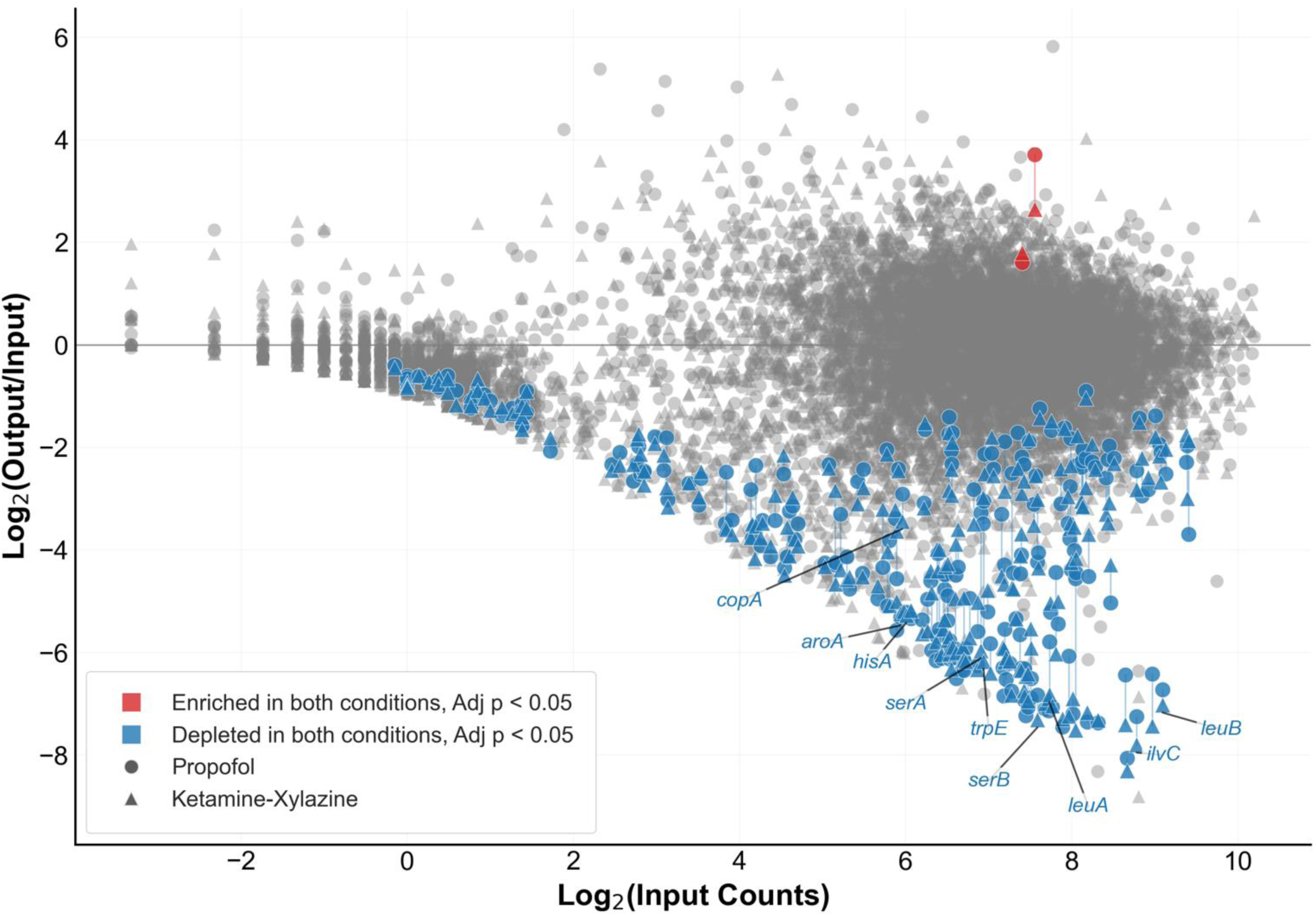
Transposon insertion sequencing results for all genes following intranasal infection in propofol-sedated mice and ketamine/xylazine-sedated mice. Each *K. pneumoniae* gene that exhibited transposon insertions in our input pool (as defined in our analysis) is represented twice in the plot, showing the performance of mutants in the gene during infection in propofol-sedated mice vs input (circles) and in ketamine/xylazine-sedated mice vs input (triangles). These data are plotted against the the representation of those mutants in the input pool. Genes that are to the right of the graph have a higher representation of mutants in the input pool and tend to produce more robust data in the INSeq analysis (e.g., larger genes with more available sites for transposon insertion, and genes for which transposon insertion does not impair growth in culture). Genes for which mutants showed a significant depletion (blue, n=217) or enrichment (red, n=2) under both sedatives are highlighted. Select genes are labeled, with a full listing of the highlighted genes listed in Tables S3 and S4, respectively.

**Figure 4.**
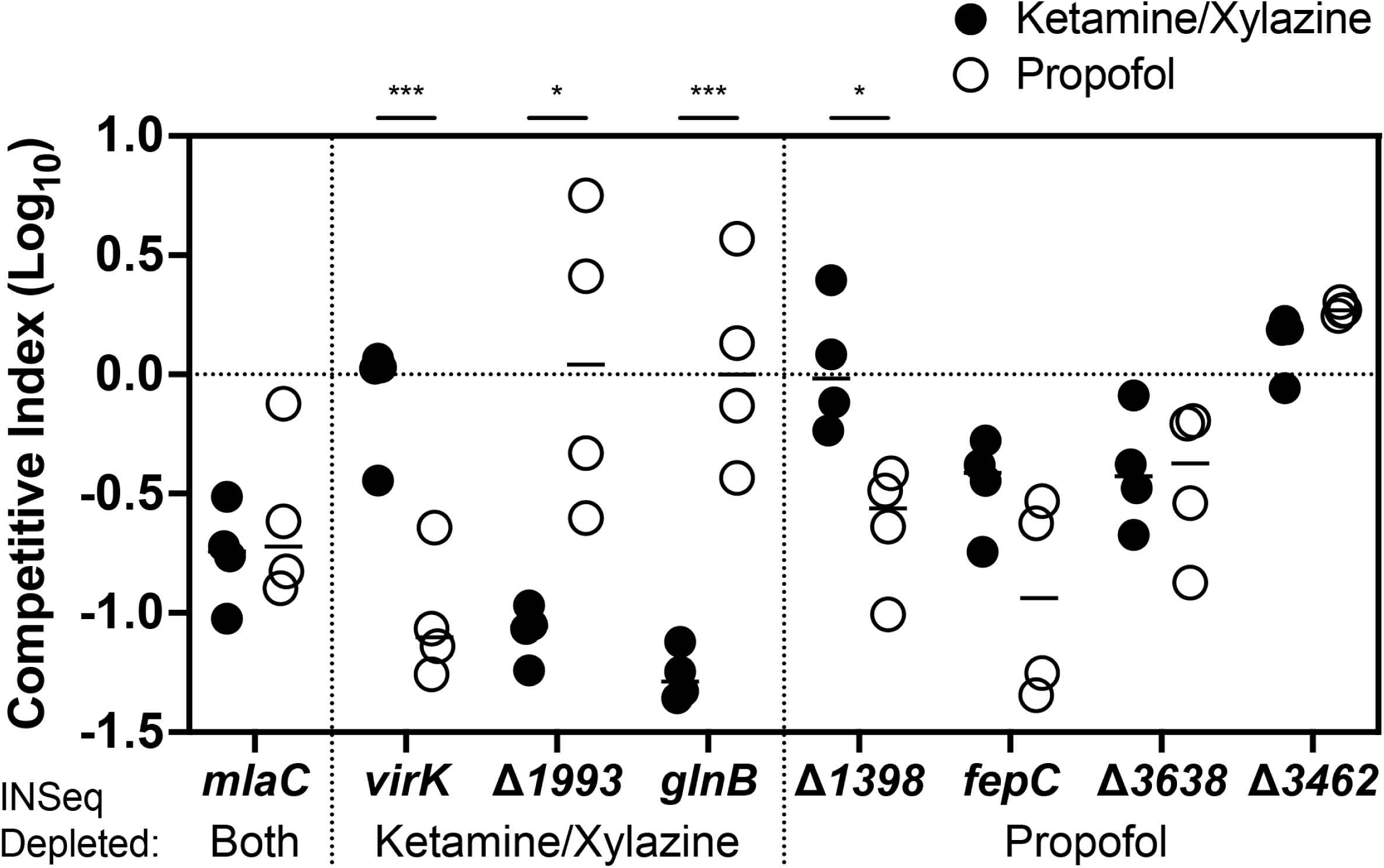
Competitive infections with in-frame deletion mutants. Competitive indices of each mutant against WT 24 hours post-infection (n=4 per group). Mice were infected with approximately 5 x 10^3^ CFU of WT and a mutant strain that were premixed and administered via intranasal route for a total 1 x 10^4^ CFU dose. Statistics on lung titers were calculated using an unpaired, two-tailed Student’s T-test. As noted in the manuscript text, depletion under INSeq was statistically significant under both sedatives for *mlaC* but was not significant for mutants under individual sedatives; the INSeq designations shown are based on the trends in Figure S1.

### Identification of *K. pneumoniae* mutants that exhibit sedative-dependent responses

We next asked whether the INSeq data had resolution to identify genes that exhibited differential responses to the two sedatives. Examination of **Figure 3** revealed variation in the response to the sedatives, even in mutants that were abundant in the input counts. A direct analysis of mutants recovered from the two output conditions (i.e., propofol-anesthetized mice compared to ketamine/xylazine-anesthetized mice) did not yield genes with significant enrichment or depletion when the TRANSIT statistical test was applied. We reasoned that this was due to the variation among transposon counts from the animal-recovered samples—in contrast to the analysis in **Figure 3** in which variation from the mouse output treatment was compared to the largely invariant input control. Therefore, we scanned the INSeq data for genes that exhibited greater depletion under one sedative. Mutants in the genes *virK* (VK055_0094/RS00455)*, VK055_1993 (RS09970)*, and *glnB* (VK055_4623/RS23395) were more substantially depleted following ketamine/xylazine sedation, while the genes *VK055_1398, VK055_3638 (RS18420), VK055_3462 (RS17525)*, and *fepC* (VK055_1930/ RS09665) were likely to be more important following propofol sedation (**Figure S1**). Additionally, we chose to investigate *mlaC (VK055_3875)* as it was identified as having a strong defect regardless of sedative choice and was representative of most of the *mla* genes that encode a phospholipid transport system (**Figure S3**).

To assess whether our approach could identify bona fide sedative-dependent mutants, we generated deletion mutants and tested each individually in competition with the wild-type parent. Competition infections were performed using a 1:1 mixture of KPPR1 and single deletion mutants. Some of the gene candidates exhibited sedative-dependent phenotypes consistent with the trends in the original transposon mutant selection (**Figure S1**). This was observed for the *ΔglnB and ΔVK055_1993* mutants that exhibited competitive defects only in the ketamine/xylazine treatment, and for *ΔVK055_1398* that demonstrated a defect only in propofol. The Δ*mlaC* mutant exhibited a competition defect in both conditions as predicted. Trends for the Δ*fepC* mutant also matched the INSeq data with slight defects under both conditions and more pronounced effect in the propofol-sedated host (**Figure S1**). However, other mutants exhibited phenotypes in the direct competition assays that differed from the trends observed in the INSeq data. The *ΔVK055_3462* mutant did not exhibit a direct competition phenotype in either condition. The *ΔVK055_3638* mutant yielded a mild and shared competitive defect in both sedatives. Unexpectedly, the Δ*virK* mutant exhibited a propofol defect in the direct competition, in contrast to the ketamine/xylazine defect in the INSeq pooled competition experiment (**Figure S1**). Although distinct from the result from INSeq, this outcome provides an additional mutant to use as a probe for host function under propofol. In summary, despite weak support for sedative-dependent mutant behavior in the INSeq data set, approximately half of the mutants that we examined validated with expected phenotypes and provide useful tools to probe the differential anesthesia response.

### Conserved virulence factors can exhibit sedative-dependent responses during mono-infection

We next wished to ask whether genes that were broadly required for *K. pneumoniae* colonization were similarly required under propofol sedation as under ketamine/xylazine. For this question, we focused on the *mlaC* mutant described above. We took advantage of the known detergent sensitivity of *mla* mutants in *E. coli* (48) to further characterize the Δ*mlaC* mutant. The *K. pneumoniae* Δ*mlaC* mutant was found to exhibit sensitivity to SDS and EDTA, a phenotype that could be reversed upon complementation with the wild-type *mlaC* expressed at the chromosomal *att*Tn*7* site (**Figure 5A, Figure S2**).

**Figure 5.**
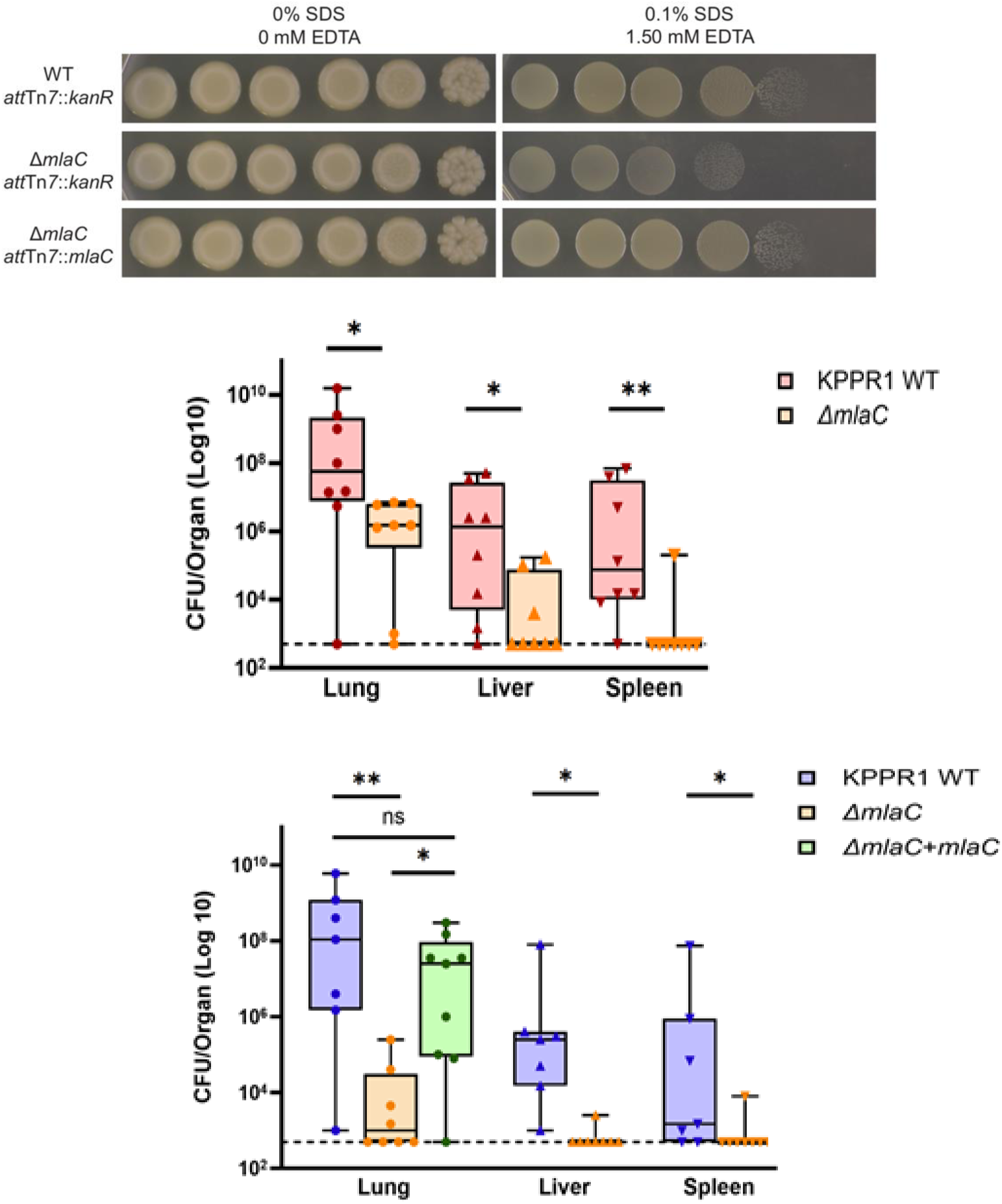
*mlaC* mutant demonstrates defects in a sedative-dependent manner in monoculture infections. (A) Saturated cultures were serially diluted and spotted onto specified media types, then grown at 37°C overnight. The growth defect of the *mlaC* mutant on SDS/EDTA is able to be complemented. (B/C) Female Swiss Webster mice (n=8-9 per group) were ketamine/xylazine sedated or propofol-sedated and infected with 2500 CFU (ketamine/xylazine sedated mice) or 250 CFU (propofol sedated mice) intranasally. The increased inoculum size for ketamine/xylazine sedated animals was necessary as mice cleared the infections at the lower (250 CFU) inoculum dose. Organs were harvested at 48 h post-infection for determination of bacterial CFU. Statistics were calculated using a two-tailed Mann-Whitney U test (*, p ≤ 0.05; **, p ≤ 0.01) and significance is compared to KPPR1 under the same sedative.

Given that propofol sedation appeared to significantly enhance mouse susceptibility to *K. pneumoniae* infection, we identified an infectious dose for propofol sedated animals that enabled mouse survival beyond 48 hours so as to provide a more extended time window for assessment of potential mutant phenotypes. An intranasal dose of 250 CFU of KPPR1 was found to establish robust and reproducible levels of infection despite being 10 to 100-fold lower than doses routinely reported in the literature (**Figure 5**) (49–51). This dose was optimized for intravenous propofol sedation as mice infected with a 10-fold higher inoculum at 2,500 CFU could not meet humane criteria for the study duration of 48 hours. For the comparison of propofol sedated animals with single Δ*mlaC* infections under ketamine sedation, a ten-fold increase to the inoculum was required for animals sedated with ketamine to detect any difference between attenuated mutant strains and WT as animals rapidly cleared lower doses of WT *K. pneumoniae* (**Figure 5**). The requirement for *mlaC* was validated for mono-infections under both sedative conditions and could be fully complemented by the introduction of *mlaC* at a neutral chromosomal locus (**Figure 5, S2**). While the requirement for a 10-fold lower CFU inoculum in propofol-sedated animals (given the increased susceptibility of these animals to *K. pneumoniae* infection) complicates a direct comparison with ketamine/xylazine sedated mouse infections, the Δ*mlaC* mutant is clearly highly attenuated in propofol-sedated mice with lung burdens comparable to ketamine-sedated controls (**Figure 5**). For example, mice infected with WT *K. pneumoniae* following propofol or ketamine/xylazine sedation exhibited comparable numbers of bacterial CFU in the lung despite the 10-fold lower dose following ketamine sedation. While the WT bacterial numbers in the lung are comparable for both groups, the Δ*mlaC* mutant is recovered in numbers that are 100-1000 fold lower following propofol sedation (**Figure 5**). Overall, our results support the requirement for specific *K*. *pneumoniae* virulence factors to establish lung infections regardless of the type of sedative used during mouse intranasal inoculation.

## DISCUSSION

Previous work identified the significant impact of brief propofol exposure on the ability of an infected host to combat bacterial bloodstream infections with either *L. monocytogenes* or *S. aureus*, both gram-positive bacterial pathogens (25, 26). Here, we have demonstrated that propofol also increases a murine host’s susceptibility to both a new route of infection (intranasal) and to a gram-negative pathogen, *K. pneumoniae*. Propofol sedation increased lung burdens approximately 100-fold compared to ketamine/xylazine-sedated mice and furthermore increased dissemination to secondary organs approximately 1,000-fold at 48 hours post-infection (**Figure 1**). We further identified that propofol sedation resulted in significantly larger areas of denuded or destroyed lung tissue (**Figure 2**), likely forming a replicative niche for rapidly dividing *K. pneumoniae in vivo* and possibly accounting for the increased lung burdens measured at 48 hours post-infection. Serious disseminated infections also resulted from a low dose inoculum (200 CFU) indicating the increased susceptibility of propofol sedated mice to *K. pneumoniae* lung infection (**Figure 5**). These data clearly demonstrate that propofol negatively impacts the host’s ability to address *K. pneumoniae* lung infection and subsequent bacteremia. Whether these changes in host immune responses resemble those previously reported for bacterial bloodstream infections, which included dramatic reductions in the populations of mature macrophages (25, 26) has not yet been determined. Lung histology indicates abundant immune cell infiltration into the infected lungs with evident differences in the resultant pathology (**Figure 2**).

Genome-wide mutagenic approaches such as signature-tagged mutagenesis (STM) and transposon mutagenesis have previously been applied as *K. pneumoniae in vivo* virulence screens and have identified dozens of putative virulence factors, which have been reviewed by Paczosa and Mecsas (30). The screen described herein yielded novel gene products by considering sedative exposure as a key variable in subsequent lung outgrowth. By comparing outputs from infected lungs to their inputs, we were able to quantify the relative requirement of numerous genes following either ketamine/xylazine or propofol sedation. This approach enabled us to identify putative novel virulence determinants and begin to define the broad impact that sedation has on pathogen fitness toward lung colonization. The choice of ketamine/xylazine for comparison was based both on the common use of this anesthetic/sedative in animal experiments, and because the drugs were previously shown to have minimal effects on the control of bloodstream bacterial infections (25, 26). Interestingly, clear effects on host infection were also evident following ketamine/xylazine sedation, suggesting that sedative use in general may impact host immunity, albeit perhaps to different degrees. This finding is consistent with a recent report from Gage et al indicating that ketamine sedation negatively impacted bacterial clearance and lung tissue pathology using a *Pseudomonas aeruginosa* pneumonia model (52).

Based on the results of the INSeq analysis and previously reported contributions of the *mla* locus to bacterial virulence (2), the *K. pneumoniae mlaC* mutant was selected for further characterization. The Mla (maintenance of outer membrane lipid asymmetry) pathway has been well characterized in *E. coli* (48). Mla comprises a transport system that shuttles phospholipids from the outer membrane to the inner membrane (48). MlaFEDB forms an ABC transporter, while MlaA is a lipoprotein attached to the outer membrane and MlaC is the periplasmic shuttle between the two. Loss of *mlaC* resulted in a significant defect under both sedative conditions (**Figure 5**). A direct comparison between propofol and ketamine/xylazine sedated animals for mutant phenotypes was somewhat constrained by the need to use increased CFU to establish infection in ketamine/xylazine sedated mice. Virulence defects similar to those observed for the *mlaC* mutant would be predicted for *mlaA*, *mlaD*, *mlaE*, and *mlaF* based on their representation in the INSeq screen (**Figure S3**). In contrast, the loss of *mlaB* does not appear to carry a significant defect based on the INSeq results, a reasonable finding considering that MlaB is a minor subunit in the transporter. *mlaA* has been recently confirmed as a virulence requirement for *K. pneumoniae* lung infections in mice (53). The perturbation of membrane lipid homeostatic mechanism is therefore likely a critical factor in the resistance of *K. pneumoniae* to host antimicrobial strategies, and this host defense may be somehow enhanced by propofol exposure, resulting in enhanced clearance of *mlaC* mutants.

Our dataset suggests numerous genes that are potentially differentially required depending on host anesthetic exposure. Further validation and characterization of these gene products is needed to provide clarity to these findings, and an expansion of the INSeq approach in the anesthetized animals would likely provide valuable additional candidates to pursue. This prediction is based on our finding of multiple interesting hits from our preliminary that were validated despite the INSeq approach not being sufficiently powered to reveal global gene requirements in propofol- versus ketamine/xylazine-sedated animals. Our results support a growing body of evidence that the choice of sedative used for anesthesia can profoundly influence the course of bacterial infection, our work goes on to show that the virulence gene repertoire required for infection may differ dependent upon sedative choice. This finding has important implications for the selection of potential new drug targets for *K. pneumoniae* infections; the most effective targets are likely to be those gene products that are required regardless of the type of sedation that may be used. Taken together, these results reveal that propofol sedation leads to the exacerbation of lung infection in a murine model of experimental pneumonia caused by the bacterial pathogen *K. pneumoniae.* Future studies aimed towards identifying colonization and virulence factors required under multiple sedation conditions should clarify which aspects of host immunity are negatively impacted by sedative exposure.

## ACKNOWLEDGMENTS

We thank Alan Hauser and Nengding (Julie) Wang for plasmid pJNW684 and use of the Promega Maxwell instrument, Jason Peters for pJMP1185 and advice on modifying the plasmid, and Emanuel Burgos for computational assistance. We thank Mercy Obarewon for assistance with histological analyses. This work was supported by the Chicago Biomedical Consortium Catalyst Award (to M.J.M. and N.E.F.), NIH grant R21AI166098 (to N.E.F. and M.J.M.), and NIH grant R35GM148385 (to M.J.M.). Its contents are solely the responsibility of the authors and do not necessarily represent the official views of the funding source.

## Supplemental Tables S1-S5

- **Table S1. NCBI SRA Accession numbers.** Data deposition for the INSeq samples.
- **Table S2. INSeq data analysis for all samples.** Shown are counts-per-million (cpm) per gene for each sample from the pyinseq pipeline.
- **Table S3. Shared depleted mutants.** Genes for which mutants showed a significant depletion under both sedatives, from the TRANSIT pipeline.
- **Table S4. Shared enriched mutants.** Genes for which mutants showed a significant enrichment under both sedatives, from the TRANSIT pipeline.
- **Table S5. Primer list.** Primers used for <u>P</u>CR during construction (P), or for <u>v</u>erification by PCR and/or Sanger sequencing to verify insertion/orientation.

