## Supplementary Figures S1-S4 for "Sedative choice alters *Klebsiella pneumoniae* lung pathogenesis and dissemination"

**Supplemental figures for ‘Sedative choice alters *Klebsiella pneumoniae* lung pathogenesis and dissemination’ by Mains et al.**

### Figure S1

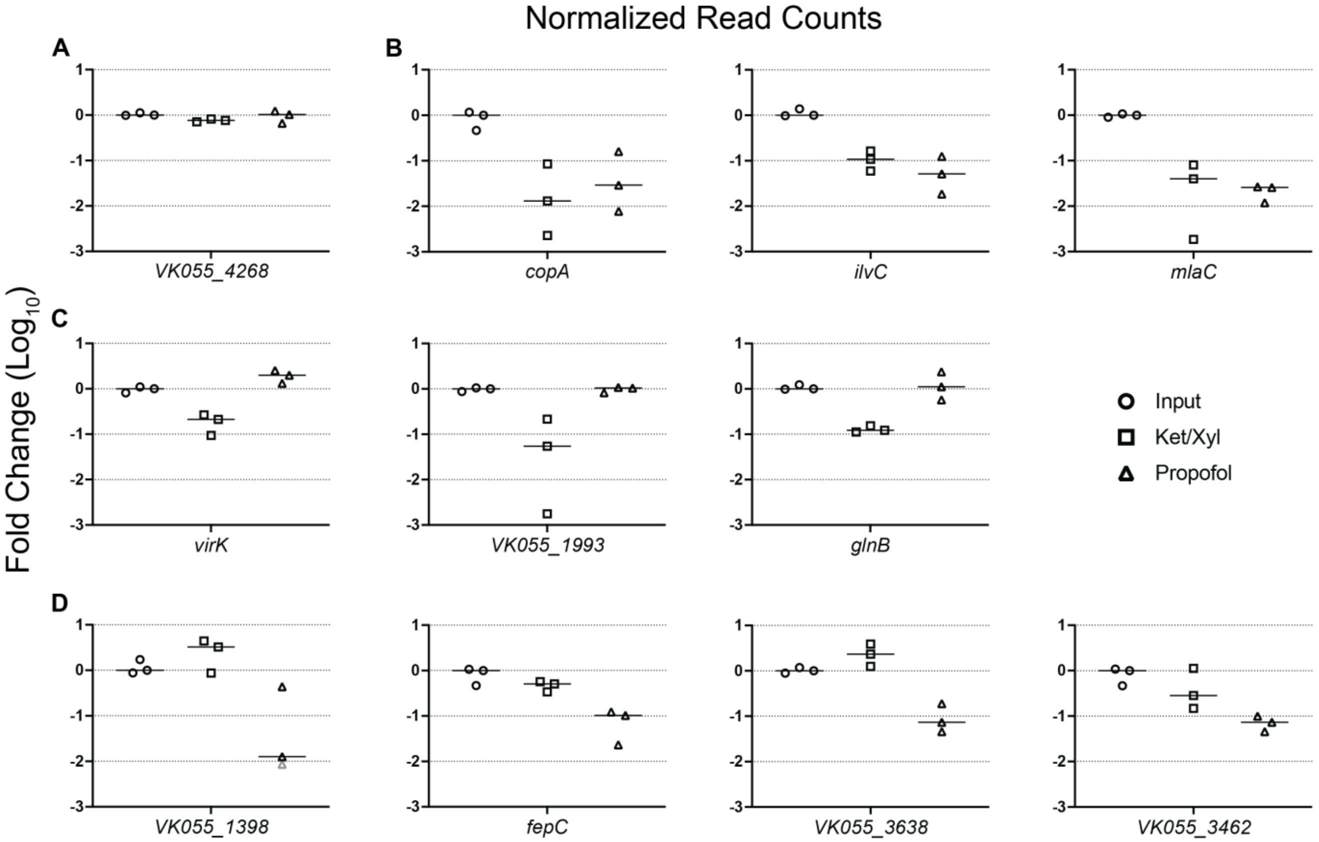


**Figure S1. Potential competitive defects identified by INSeq.** Competitive results for the input (circles), control-treated mice (squares), and propofol-treated mice (triangles). The values are normalized to the median of the three inputs. A) The first graph contains an example of a gene not relevant for virulence (*VK055_4268*) in either sedative condition. B) The remaining three graphs in the first row show genes with defects regardless of sedative used. Genes *copA* and *ilvC* were previously validated. *mlaC* is a general virulence factor. C) The second row (*virK*, *VK055_1993*, *glnB*) are genes for which mutants exhibited defects in ketamine-sedated mice only. D) Third row (*VK055_1398*, *fepC*, *VK055_3638*, *VK055_3462*) are genes for which mutants exhibited defects in propofol-sedated mice. Points in gray had zero normalized read counts and were set to 0.1 for the purpose of calculating log fold change. Data are from the pyinseq analysis.

Figure S2

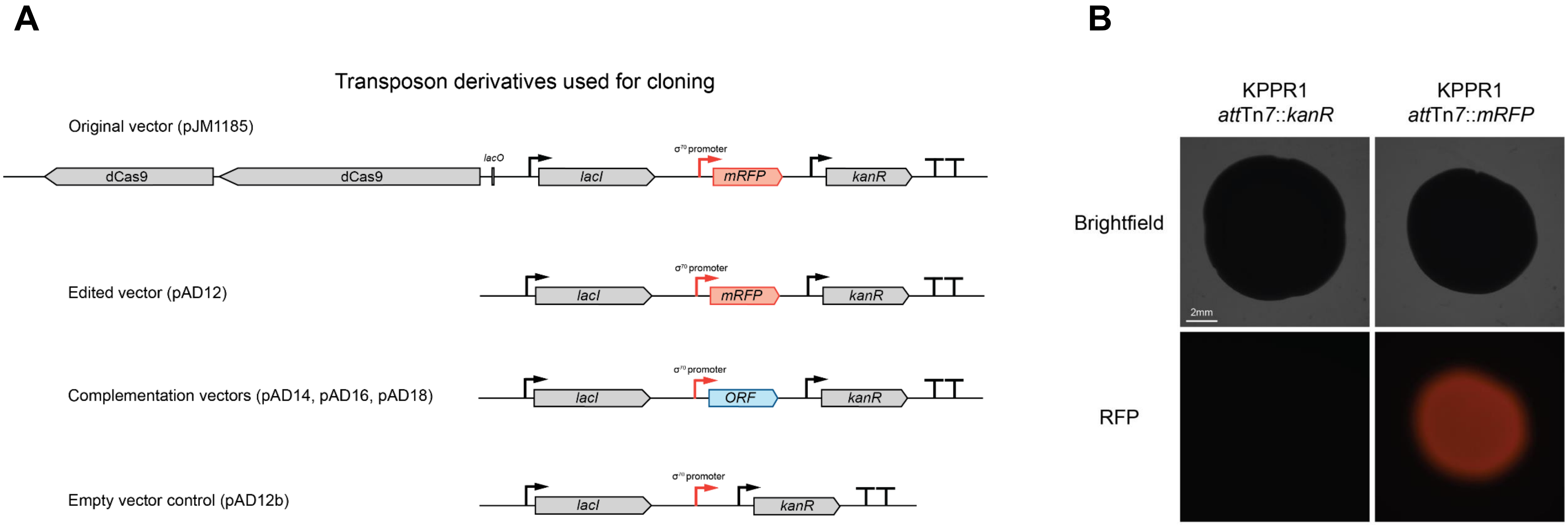


**Figure S2. Tn7 complementation approach.** (A) The *cas9* gene was removed from pJM1185 to create plasmid pAD12. Since this plasmid expresses *mRFP*, we used it to test expression levels in KPPR1 by confirming red fluorescence as seen in panel B. To create complemented strains, the genes were cloned in to replace the *mRFP* gene creating plasmids pAD14, pAD16, and pAD18 depending on the gene inserted. To create an empty vector control, *mRFP* was removed to create plasmid pAD12b. (B) From overnight growth in LB, 8 µl culture was spotted onto LB and grown for 24 h at 30 °C before imaging.

Figure S3

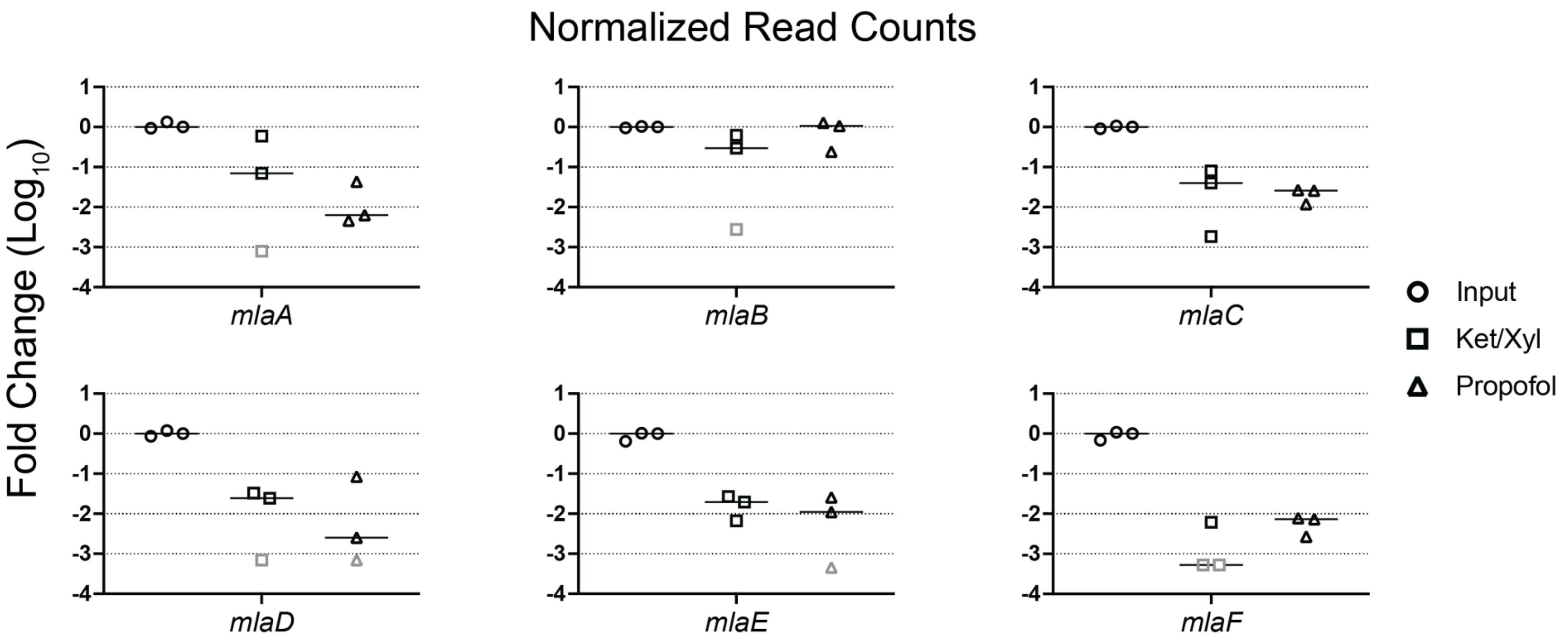


**Figure S3. Potential *mla* operon defects identified by INSeq.** Competitive results for the input (circles), control-treated mice (squares), and propofol-treated mice (triangles). The values are normalized to the median of the three inputs. Points in gray had zero normalized read counts and were set to 0.1 for the purpose of calculating log fold change.

Figure S4

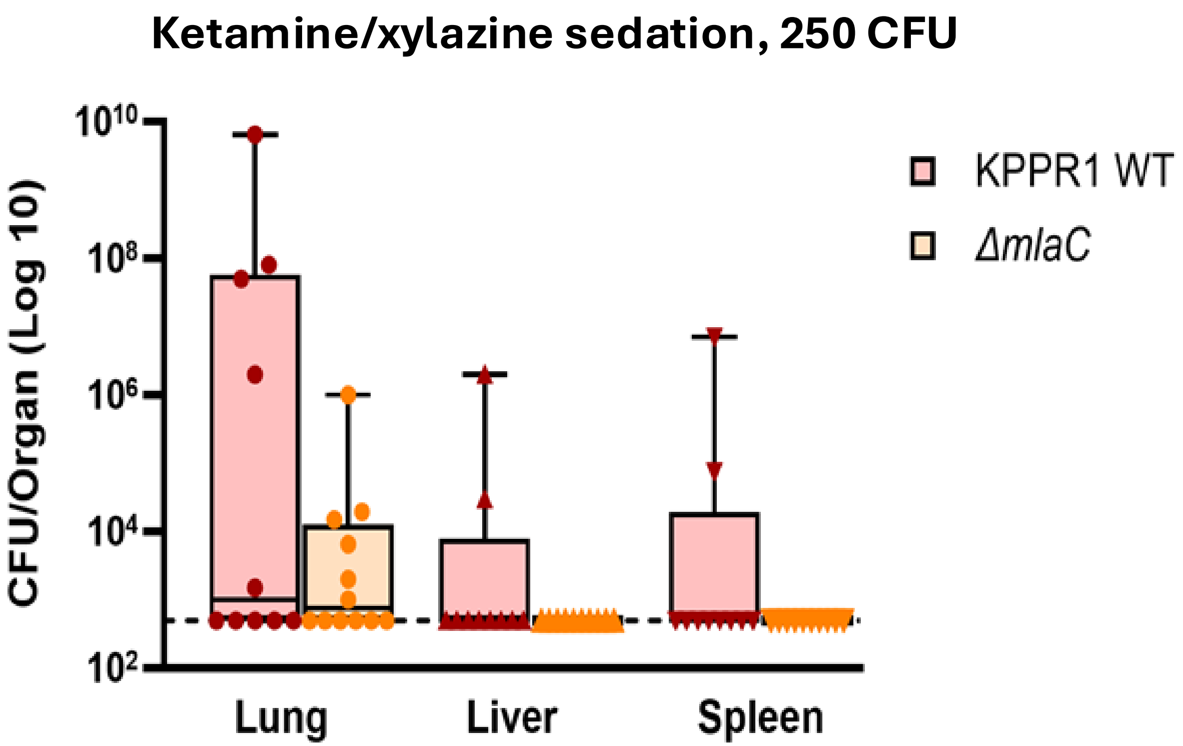


**Figure S4. Mice sedated with ketamine/xylazine are less susceptible to low dose KPPR1 infection.** Female Swiss Webster mice were sedated with ketamine/xylazine and infected with 250 CFU KPPR1.Organs were harvested at 48 h post-infection to assess bacterial CFU. At the low infectious dose of 250 CFU the median values are near the level of detection for mice infected with either WT or Δ*mlaC* .
